# A spatially-aware unsupervised pipeline to identify co-methylation regions in DNA methylation data

**DOI:** 10.1101/2025.11.13.688356

**Authors:** Siddhant Meshram, Arindam Fadikar, Ganesan Arunkumar, Suvo Chatterjee

## Abstract

DNA methylation (DNAm) plays a central role in modern epigenetic research; however, the high dimensionality of DNAm data comprising hundreds of thousands of spatially ordered probes continues to present major analytical challenges. The multiple testing burden in these data introduces redundancy and reduces statistical power, contributing to the limited reproducibility often observed in association studies. Moreover, DNAm probes frequently exhibit correlated methylation patterns with neighboring sites, reflecting underlying biological co-regulation and spatial dependence along the genome. Ignoring these spatial correlations can bias parameter and standard error estimates, inflate type I error rates, and obscure biologically meaningful effects. Existing methods for detecting methylation co-regulation and reducing DNAm data dimensions, typically rely on fixed distance or correlation thresholds and arbitrary hyperparameter settings that lack data adaptivity. In this study, we introduce SACOMA (Spatially-Aware Clustering for Co-Methylation Analysis), a flexible, data-driven, and unsupervised framework designed to identify co-methylated regions which are genomic regions where adjacent sites show correlated methylation levels. SACOMA employs spatially constrained hierarchical clustering to group neighboring DNAm sites based on both spatial proximity and methylation similarity. A tunable, data-adaptive mixing parameter allows SACOMA to avoid rigid assumptions and remain robust to hyperparameter choices. Although developed for DNAm array data, SACOMA provides a generalizable framework applicable to any data exhibiting spatial dependence, enabling the identification of spatially correlated features across diverse domains. Through extensive simulations, SACOMA demonstrated superior sensitivity while maintaining effective false-positive control compared to existing methods. In population-level DNAm data analyses, SACOMA successfully identified biologically relevant co-regulated methylation regions with functional roles. Overall, SACOMA reduces the multiple-testing burden and enhances both the discovery and specificity of statistical associations, leading to improved reproducibility and more reliable biological inference.

## 1 Introduction

DNA methylation (DNAm), the covalent addition of a methyl group to the fifth carbon of cytosine residues in the context of cytosine-phosphate-guanine (CpG) dinucleotides, is one of the most studied and well understood epigenetic modifications in the human genome [1, 2]. This biochemical mark plays a pivotal role in a wide range of biological processes, including the establishment and maintenance of cellular identity, regulation of gene expression, inactivation of the X-chromosome, genomic imprinting and suppression of transposable elements [3, 4]. Aberrations in DNA methylation patterns are hallmarks of many diseases, particularly cancers, where global hypomethylation and region-specific hypermethylation disrupt normal gene regulation and promote tumorigenesis [5]. Moreover, recent studies have underscored the dynamic nature of the methylome, highlighting how DNA methylation responds to environmental exposures such as tobacco smoke, air pollution, diet and psychosocial stress, thus serving as a molecular integrator of gene-environment interactions [6, 7]. This combination of biological importance, disease relevance and environmental responsiveness has placed DNA methylation as a critical target in efforts to understand the etiology of complex traits and as a promising biomarker for precision medicine.

High-throughput platforms such as the Illumina Infinium 450K and EPIC BeadChip have enabled researchers to profile DNA methylation at hundreds of thousands of CpG sites simultaneously, covering regions enriched for gene promoters, CpG islands and regulatory elements [8, 9]. Once these data are generated, investigators typically conduct epigenome-wide association studies (EWAS) to link differential methylation patterns to phenotypes, traits, exposures or disease outcomes. EWAS commonly rely on single-site models that test each CpG independently. Although this approach is straightforward and provides high resolution information, it suffers from severe multiple testing penalties given the vast number of sites examined, often resulting in limited power to detect modest yet coordinated methylation changes that could be biologically meaningful [10]. Recognizing these limitations, in the recent years there has been a methodological shift towards region-based analyses, which aggregate information across contiguous CpG sites to identify differentially methylated regions (DMRs). Such region-level investigations not only address the multiple testing burden but also align more closely with the biological reality that regulatory methylation changes frequently manifest across groups of neighboring CpGs rather than isolated single sites [5, 11].

Currently, region-based analysis in DNAm array datasets can be broadly categorized into two methodological frameworks, supervised and unsupervised. Supervised frameworks first identify CpG sites that show significant associations with a phenotype or exposure and then combine adjacent significant sites into co-methylated regions which depict differential methylation also called as DMRs. Majority of these tools employs its own strategy to aggregate nearby CpGs typically using genomic proximity and p-value significance thresholds to define contiguous regions of coordinated change [12, 13]. This phenotype-guided approach provides a practical and computationally efficient framework, particularly when strong site-level signals exist. However, their dependence on initial single-site testing limits their ability to detect subtle coordinated methylation changes that are distributed across several CpGs. Such changes may not reach site-level significance and therefore remain undetected during region formation. In contrast, unsupervised methods first define co-methylated regions independently of any phenotype information by relying solely on intrinsic methylation correlation patterns and genomic distance among CpG sites, and subsequently perform association analysis. These models aim to uncover the natural structural organization of the methylome, reflecting the underlying chromatin and regulatory landscapes without bias from the outcome of interest. Although unsupervised models avoid the limitations of supervised approaches, they can still face challenges when they fail to accurately capture true co-methylation structure either by grouping weakly correlated CpGs or fragmenting biologically coherent regions. The resulting regions can introduce noise, inflate false positives or reduce statistical power in downstream region-based testing.

A crucial part of unsupervised region-based methylation analysis is the process of identifying co-methylated regions, wherein methylation levels of neighboring CpG sites along the genome tend to be correlated [14, 2]. This spatial dependency reflects the fact that CpGs situated within the same chromatin context such as promoters, enhancers, CpG islands or gene bodies are often subject to common regulatory influences [4]. Chromatin accessibility, nucleosome positioning, histone modifications, and transcription factor binding collectively shape local methylation environments, resulting in coordinated methylation dynamics across short genomic intervals [15, 4]. Such coordination is biologically meaningful, because it often mirrors the functional states of the underlying genomic region. From a statistical standpoint, co-methylation implies a dependency structure that violates the assumption of independence across CpGs, an issue that must be considered when designing association studies. By explicitly modeling these dependencies, region-level approaches can capture subtle but consistent methylation changes distributed between groups of CpGs. As a result, they offer improved power, greater robustness to noise and more biologically meaningful results compared to site-level analyses. Importantly, by focusing on co-methylated domains rather than individual CpGs, these models facilitate the detection of coordinated regulatory mechanisms and provide insights into the spatial organization of the epigenome.

In the past decade, some models have been developed for identifying co-methylated regions such as Aclust2 (updated version of Aclust) [16] [17] and coMethDMR [18]. Although these models have been the predominant choices for unsupervised, region-based analysis, they exhibit notable shortcomings and methodological gaps, particularly in accurately identifying co-methylated regions, which represents a crucial step in these analytical pipelines. For instance, Aclust/Aclust2, uses a fixed distance threshold to define co-methylated regions based on the assumption that CpGs within this threshold exhibit similar methylation behavior, an assumption that may not universally hold across different genomic contexts. This arbitrary cutoff can split biologically coherent regions, thereby reducing statistical power in sparse regions and making the model more sensitive to local CpG density and correlation structure rather than true biological co-methylation patterns. Similarly, coMethDMR identifies co-methylated sub-regions by computing pairwise correlations (rdrop values) among CpGs within predefined regions, retaining only those with rdrop > 0.4. However, this fixed global threshold is arbitrary and can substantially influence results as small changes can alter the number and size of detected regions, merging weakly correlated CpGs and increasing false positives or excluding moderately correlated yet biologically relevant ones and increasing false negatives. Because it applies the same cutoff across all genomic contexts, it fails to account for differences in CpG density and correlation structure, reducing adaptability, reproducibility across datasets, and biasing detection toward regions with uniformly strong correlations while overlooking partially co-methylated domains.

In this study, we introduce SACOMA (<u>S</u>patially-<u>A</u>ware Clustering for <u>Co</u>-<u>M</u>ethylation <u>A</u>nalysis), a flexible and data-adaptive unsupervised framework for identifying co-methylated regions that enhances the crucial first step of unsupervised region-based methylation analysis. The primary objective of this study is the optimization of identifying co-methylated regions using an unsupervised approach, ensuring that subsequent analyses whether simple or complex are based on more accurate and biologically coherent methylation clusters. SACOMA adaptively learns CpG groupings by jointly optimizing spatial proximity and methylation homogeneity through a tunable mixing parameter, requiring minimal user-defined inputs and avoiding the rigid distance constraints and sensitivity to tuning parameters. Through comprehensive benchmarking, SACOMA demonstrated an optimal balance between sensitivity and specificity while maintaining computational efficiency compared to popular unsupervised tools. Beyond simulations, co-methylated regions identified by SACOMA in population-level DNAm array datasets were biologically relevant as these regions were known to exhibit coordinated methylation regulation and have functional relevance. This concordance between simulation and empirical results underscores SACOMA’s reliability, scalability, and biological validity. Collectively, these findings highlight that SACOMA exhibits superior performance in correctly identifying co-methylated regions, providing a powerful and biologically informed foundation for downstream region-based methylation analyses.

## 2 Materials and methods

Identifying co-methylated regions can be formulated as a problem of spatially constrained clustering, where the goal is to detect groups of CpG sites that exhibit both high methylation correlation and spatial contiguity within the genome. In essence, this involves clustering CpGs based on their methylation profiles while simultaneously enforcing a proximity constraint defined by genomic distance in base pair thresholds. Statistically, this dual criterion reflects the biological reality that DNA methylation is often regionally coordinated rather than randomly distributed across distant loci. Traditional clustering methods typically overlook spatial autocorrelation, the intrinsic property that nearby CpG sites tend to have more similar methylation levels than distant ones [19]. Spatially constrained clustering addresses this by incorporating spatial relationships through neighborhood structures, distance metrics, or spatial weight matrices, as demonstrated in methods such as DBSCAN [20] and its temporal extension ST-DBSCAN [21], or spatially constrained hierarchical clustering [22] and GeoSOM [23]. However, these approaches often suffer from rigid spatial constraints or sensitivity to local connectivity. The ClustGeo algorithm [24] overcomes these limitations by introducing a convex combination of two dissimilarity matrices, one representing methylation similarity and another representing spatial proximity, thus enabling a soft, tunable integration of spatial information while preserving the interpretability and hierarchical nature of clustering. Building on this foundation, we develop SACOMA to robustly identify co-methylated genomic regions by jointly modeling methylation correlation and spatial organization in spatially constrained hierarchical clustering framework, thereby capturing biologically meaningful epigenetic domains.

Specifically, SACOMA’s framework starts by creating genomic bins that group adjacent CpG sites into candidate regions based on their genomic proximity and density. The Illumina array manifest file, which contains the genomic coordinates and probe annotations for each CpG site, is first divided by chromosome and ordered by genomic position to preserve spatial continuity across loci. CpG sites that lie within a user-defined maximum distance are then merged into the same genomic bin, forming contiguous CpG clusters that represent potential co-methylated regions. Clusters containing at least *n* number of CpGs are retained to ensure sufficient CpG density and biological relevance. For each resulting bin containing *n* CpG sites, the procedure begins by constructing a *n × n* pairwise correlation matrix, *C* = [*ρ*_*ij*_], where *ρ*_*ij*_ denotes the correlation coefficient between the methylation profiles of sites *i* and *j* across all samples. The correlation matrix is thresholded at a user-specified level *r* ∈ [0, 1] to determine which CpG pairs exhibit sufficient co-methylation to be considered part of the same region. This results in a binary adjacency matrix *A* = [*a*_*ij*_] where *a*_*ij*_ = II(|*ρ*_*ij*_| ≥ *r*), indicating whether sites *i* and *j* are considered sufficiently correlated. This adjacency matrix is then transformed into a feature dissimilarity matrix 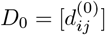 by defining 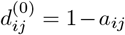. As such, *D*_0_ encodes the absence of co-methylation relationships and is used to quantify within-cluster dissimilarity in methylation levels. Simultaneously, a spatial dissimilarity matrix 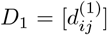 is constructed based on the genomic positions of the CpG sites. Each site is assigned a scalar genomic coordinate (e.g., midpoint between start and end probe locations), and the dissimilarities are computed as 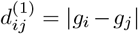, where *g*_*i*_ and *g*_*j*_ are the genomic positions of sites *i* and *j*, respectively. This matrix captures the linear genomic distance between sites and serves as a proxy for spatial structure.

To identify the optimal balance between methylation similarity and spatial proximity, SACOMA performs model selection over a predefined grid of candidate mixing parameters {*α*_1_, …, *α*_*J*_} ⊂ [0, 1]. For each *α*_*j*_, hierarchical clustering is applied to *D*_0_ and *D*_1_ by iteratively merging clusters to minimize a convex combination of within-cluster pseudo-inertias:

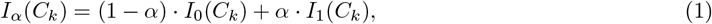

where *C*_*k*_ is the *k*-th cluster, and *I*_0_(*C*_*k*_) and *I*_1_(*C*_*k*_) represent within-cluster dispersion in the feature and spatial dissimilarity spaces, respectively. The mixing parameter *α* ∈ [0, 1] controls the trade-off between methylation homogeneity and spatial compactness.. The resulting dendrogram is cut to produce a partition 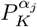 into *K* clusters, where *K* is typically set to ⌊*n/*3⌋ to encourage fine resolution. For each partition, the proportion of explained pseudo-inertia is computed in both the feature and spatial spaces:

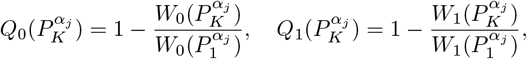

where *W*_0_(·) and *W*_1_(·) denote the total within-cluster pseudo-inertias under *D*_0_ and *D*_1_, respectively. The optimal mixing parameter is then selected as:

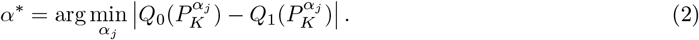

Using the selected *α*∗, a final hierarchical clustering is performed to generate a dendrogram. This dendrogram is then cut at height zero, corresponding to the lowest possible aggregation level, to extract all subclusters formed during the agglomerative process. Among these, SACOMA retains only those clusters that contain > 2 CpG sites. If no such clusters are found, the bin is classified as *noise/non-co-methylated*; otherwise, all retained clusters are labeled as *signal/co-methylated regions*. These steps are repeated independently for each genomic bin. For computational efficiency, bins can be processed in parallel. The complete workflow is formalized in algorithm 1.

### Algorithm 1

SACOMA

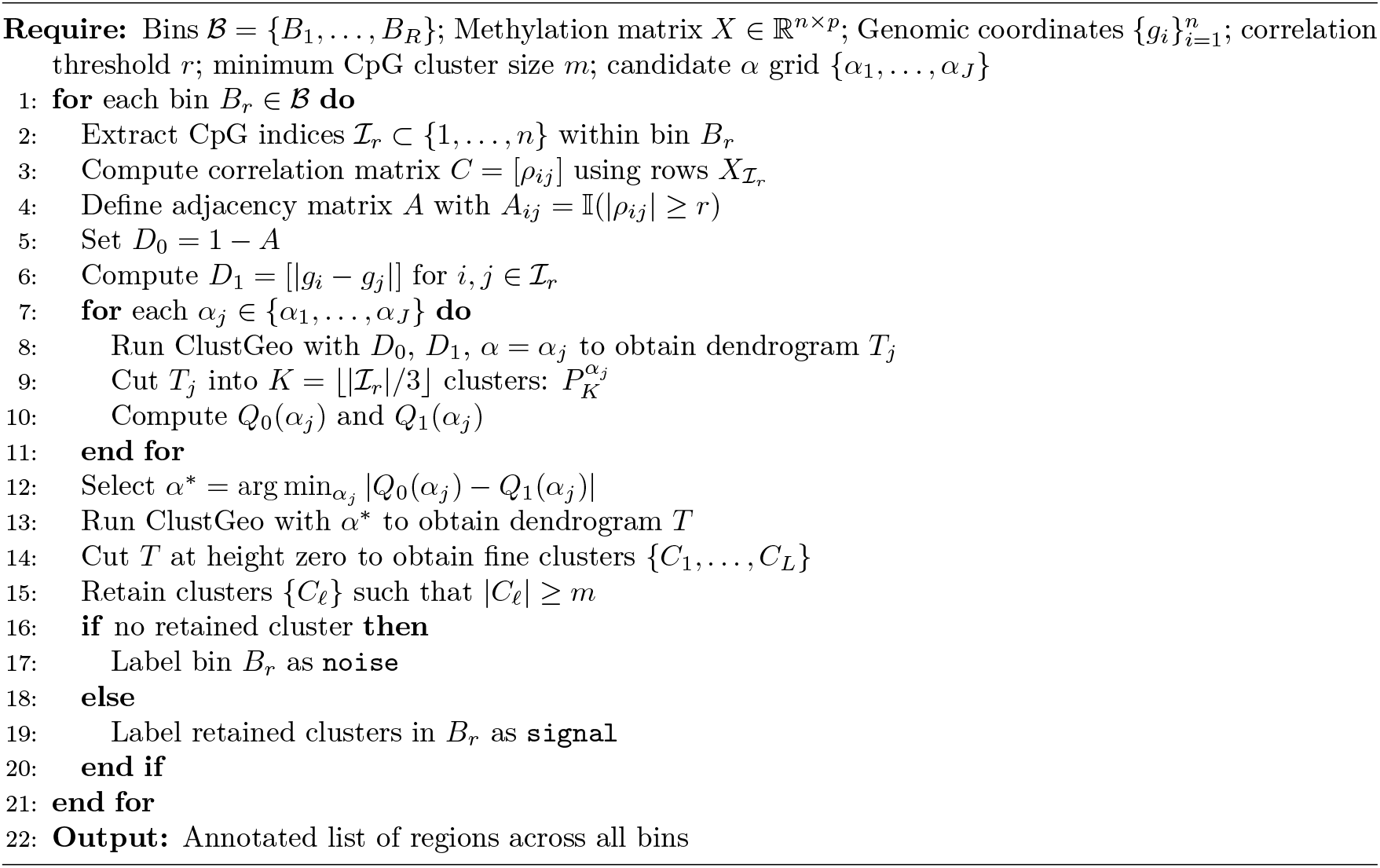

## 2.1 Statistical Models for identifying co-methylated regions

In this section, we provide an overview of the statistical methods for identifying co-methylated regions evaluated alongside SACOMA, which are coMethDMR [18] and Aclust2 [17].**coMethDMR:** coMethDMR [18] is designed to identify co-methylated regions by detecting subsets of CpGs within predefined genomic areas that show strong correlation in their methylation patterns. It first groups CpGs based on genomic annotations, then applies a filtering step to select only those contiguous CpGs that are highly correlated with each other. To do this, it uses the rdrop statistic, which measures how strongly each CpG correlates with the sum of the others in the region. Contiguous CpGs with correlation above a chosen threshold are retained, forming a co-methylated sub-region. This process ensures that only the most consistent and biologically meaningful clusters of CpGs are identified as co-methylated regions. coMethDMR is available as an open-source R package

**Aclust2.0:** Aclust2.0 [17] is an updated unsupervised framework for identifying clusters of co-methylated CpG sites from Illumina 450K, EPIC, and MM285 arrays. It builds on the original Aclust [16] model by integrating spatial proximity and correlation among adjacent CpGs to define co-methylated regions, independent of phenotype information. The pipeline automates preprocessing, clustering, and annotation by first filtering out CpG sites with excessive missing values and then grouping neighboring CpGs located within a fixed genomic distance into candidate co-methylated clusters.

### 2.2 CpG and Genic Annotation

To annotate CpGs, co-methylated regions identified by each method were extracted. The CpGs of those regions were then annotated by joining them with the array manifest file (HM450 or EPICv1). The predominant annotation per region was identified based on frequency and the proportions of regions across CpG neighborhood (Island, Shore, Shelf, OpenSea) were summarized. The genic annotations were performed in R using the annotatr [25], GenomicRanges [26] and TxDb.Hsapiens.UCSC [27, 28] packages. For each method, co-methylated regions were identified and their associated CpG sites were converted to genomic coordinates. These CpGs were annotated against gene-based features obtained from the appropriate TxDb object and annotatr gene models. CpGs located 1-5 kb upstream of transcription start sites were identified separately using promoter definitions from GenomicRanges. Each CpG was assigned to one of several functional categories (Promoters, 1-5 kb TSS, Exons, Introns, 5’UTRs, 3’UTRs or Intergenic) based on hierarchical priority rules. The annotations were summarized by assigning each region the predominant genic category among its CpGs.

### 2.3 Data Description

In this study, we obtained two publicly available methylation dataset from the Gene Expression Omnibus (GEO; https://www.ncbi.nlm.nih.gov/geo/) database under accession number GSE281199 (based on 450K) and GSE169338 (based on EPICv1). The 450K dataset originates from the EVOIMMUNOPOP project (Human Evolutionary Immunogenomics: Population Genetic Variation in Immune Responses) [29]. It contains 175 samples profiled using the Illumina’s Infinium HumanMethylation450 Beadchip, which assesses over 480,000 CpG sites throughout the genome. The raw DNAm data were color corrected and the background was subtracted using GenomeStudio software (Illumina, v2011.1). Data analysis was performed in R (v3.5.1). Pre-processing was performed with the minfi (v1.32.0) [30] and methylumi (v2.28.0) [31] packages. During quality control, a subset of probes was removed, which are SNP control probes, probes on the X and Y chromosomes, probes with detection p-values > 0.05 in more than 1% of samples, probes with missing values in more than 5% of samples, probes with limited bead count (< 3) in more than 5% of samples and polymorphic CpG probes. Samples with more than 1% missing probes were also excluded from further analysis. To adjust for probe design differences between Infinium type I and II probes, beta-mixture quantile (BMIQ) normalization [32] was performed. Batch effects associated with chip and row were corrected using ComBat function [33] of sva package [34]. DNAm levels were expressed as beta-values, representing the ratio of methylated probe intensity to total probe intensity [29].

This EPICv1 dataset investigated the pattern of variation in DNAm during pregnancy among preterm mothers in the GARBH-Ini cohort [35]. Genome-wide DNAm profiling was performed on peripheral blood DNA collected from pregnant individuals throughout gestation and at delivery. The dataset contain 34 samples with 4 timepoints each, including term and preterm deliveries. It was profiled using Illumina’s Infinium MethylationEPIC BeadChip, which measures over 850,000 CpG sites throughout the genome. Bisulfite conversion of genomic DNA was carried out using the EZ DNA Methylation Kit (Zymo Research, CA, USA) and the arrays were scanned using the iScan system (Illumina) with GenomeStudio software (v2011.1, Illumina) to obtain raw intensity data and generate beta-values. Probes with fewer than three beads in at least 5% of the samples, probes mapping to non-CpG sites, those overlapping known SNPs, multi-hit probes, and probes located on sex chromosomes were removed using the ChAMP package (v2.8.3) [36] in R (v3.6.3). Data were normalized with beta-mixture quantile normalization [32] to adjust for the two probe types present on the array. Quantile normalization was also applied to equalize the intensity distributions between samples. The technical variation associated with chip position and batch effects was corrected using ComBat [33] function in R [35]. For the analysis, samples collected at the time of delivery was used.

### 2.4 Simulation Framework and Settings

To comprehensively evaluate methods identifying co-methylated regions in DNAm array data, we generated synthetic DNAm datasets containing co-methylated regions using our simulation framework. Our simulation framework leverages existing DNAm array datasets as templates to ensure that the simulated data closely resemble real-world DNAm profiles. Two template datasets were used: one based on the 450K array and the other on the EPICv1 array. The 450K dataset (GSE281199), obtained from the EVOIMMUNOPOP project (Human Evolutionary Immunogenomics: Population Genetic Variation in Immune Responses), consisted of 175 samples [29]. The EPICv1 dataset (GSE169338) was derived from DNAm profiling of peripheral blood DNA collected from pregnant individuals throughout gestation and at delivery, comprising 34 samples with 4 timepoints, including term and preterm deliveries [35]. In our simulation context, samples collected only at the time of delivery were used. Using these two platforms allowed us to evaluate model performance across DNAm datasets of different dimensions. First, methylation beta values were loaded from the 450K or EPICv1 platform, along with the corresponding manifest files containing probe annotations. The beta value matrix was cleaned by converting all entries to numeric values and removing probes with missing data. To ensure consistency, the manifest was filtered so that only probes present in the methylation data were retained. Genomic regions were then defined by grouping CpG sites that were located within a maximum allowed genomic distance, which was set to either 200, 500 or 1000 bp. Each region contained exactly 3, 5 or 7 CpG sites. For power simulations, we assessed the similarity of methylation patterns within each candidate region by calculating pairwise correlations between CpG sites. Regions with an average correlation of at least 0.5 were retained as co-methylated regions and treated as true signals, and those failing to meet this threshold were considered noise. To balance the evaluation dataset, noise regions were randomly sampled in proportion to the number of signal regions. All regions were then combined into a single dataset, annotated with correlation statistics and labeled as signal or noise. Finally, the regions were ordered by genomic coordinates and CpG probes were grouped accordingly, yielding a dataset with biologically meaningful co-methylated regions alongside controlled noise regions. For the null simulations, we followed the same pre-processing and region-definition steps but imposed the opposite correlation filter. Instead of selecting highly correlated regions, we restricted the dataset to regions with weak correlations, defined by pairwise correlations capped at 0.4. From these, a random subset of 500 regions was sampled to construct the evaluation dataset and all CpGs were labeled as ‘noise’. This ensured that the null dataset contained no true co-methylation signal, thereby providing an appropriate baseline for method evaluation.

For power simulations, each synthetic dataset contained varying numbers of regions, defined by the number of CpGs per region and the maximum gap (maximum allowed genomic distance (in base pairs) between adjacent CpG sites for them to be grouped into the same region) permitted between CpGs within a region. We considered three types of length of regions with 3, 5 and 7 CpGs per region and three maximum gap thresholds (200 bp, 500 bp and 1000 bp). In each scenario, the numbers of co-methylated and non–co-methylated regions were kept equal. Co-methylated regions were simulated with a minimum correlation of 0.5, while non–co-methylated regions had correlations below 0.5. A complete summary of all simulation scenarios evaluated across the models is presented in **STable 1 and STable 2**. For null simulations, each dataset comprised 500 regions with correlations below 0.4, representing non–co-methylated regions. The same values for CpGs per region and maximum gap were applied as in the power simulations. Each scenario was replicated 100 times to ensure consistency and statistical reliability.

## 3 Results

### 3.1 Depiction of co-methylated regions in DNAm array data

Co-methylation refers to the correlated methylation of neighboring CpG sites across the genome, reflecting coordinated regulation of DNA methylation within local chromatin domains [18]. This spatial dependence arises because CpGs within the same genomic context such as promoters, enhancers, CpG islands or gene bodies often experience shared epigenetic influences driven by chromatin accessibility, histone modifications, and transcriptional activity [15, 4]. From a biological perspective, such coordinated methylation changes capture functionally coherent regulation of gene expression and maintenance of genome stability. Statistically, accounting for co-methylation is essential because neighboring CpGs exhibit strong correlation structures that violate the assumption of independence often made in EWAS. Aggregating correlated CpGs into regional units therefore mitigates the multiple-testing burden and enhances power to detect subtle, yet biologically meaningful, methylation changes [17, 18].

To depict the presence of co-methylated regions in array-based DNA methylation data, in this study we utilized two publicly available population-level DNAm array datasets obtained using the 450K and EPICv1 platforms (see Methods). Using the corresponding manifest files, CpG sites were grouped into regions containing at least three CpGs located within 200 base pairs of each other. For each region, we computed pairwise Spearman correlations [37] among CpGs and summarized the distribution of intra-region correlations **(Figure 1A-B)**. Both datasets exhibited a substantial proportion of regions with moderate to high within-region correlation (*ρ* > 0.4), providing strong evidence for locally co-methylated blocks of CpGs. Interestingly, regions containing a greater number of CpGs tended to show higher correlation, suggesting that CpG density and physical proximity jointly contribute to co-methylation patterns.

**Figure 1.**
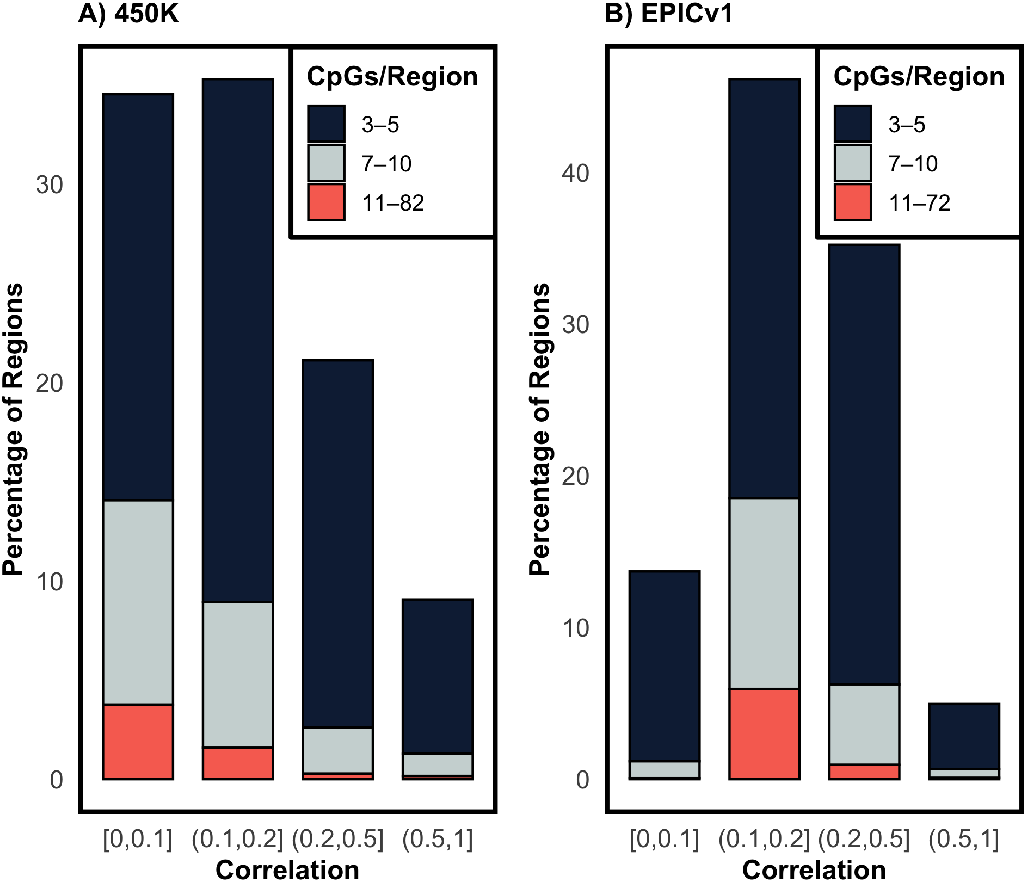
Depiction of co-methylation patterns in DNA methylation array data. Distribution of intra-region CpG correlations in 450K (panel **A**) and EPICv1 (panel **B**) datasets. Regions were defined as groups of ≥ 3 CpGs within 200 bp. The plots show the percentage of regions across correlation intervals, stratified by the number of CpGs per region.

To further characterize the genomic context of these co-methylated regions, we examined their distribution across CpG neighborhoods and genic annotations for both the 450K and EPICv1 datasets (**Table 1**). In both platforms, a notable proportion of co-methylated regions localized within CpG islands (41.7% in 450K and 31.4% in EPICv1) and promoter regions (21.6% in 450K and 23.4% in EPICv1). This enrichment is consistent with findings that CpG islands and promoters frequently exhibit strong co-methylation, as neighboring CpGs in these regulatory regions tend to share similar methylation states due to coordinated chromatin accessibility and transcriptional regulation [38, 14, 2, 4]. In total, 11.76% of the regions in the 450K dataset and 9.08% in EPICv1 were identified as co-methylated. The remaining regions constituting 88.24% and 90.92% of the genome, respectively showed little to no co-methylation correlation. Effectively filtering out such non-co-methylated regions is a crucial dimension reduction step that can substantially decrease the multiple-testing burden and improve computational efficiency. Failure to perform this reduction may inflate false positives or diminish power in downstream analyses by introducing spurious regions into testing frameworks. Therefore, a powerful and flexible model for co-methylation detection is essential to distinguish truly correlated domains from background noise. Collectively, these observations illustrate the structured nature of co-methylation across the genome and provide a clear depiction of co-methylated regions in DNAm array data, highlighting the biological and analytical importance of accurately identifying such regions.

**Table 1.**
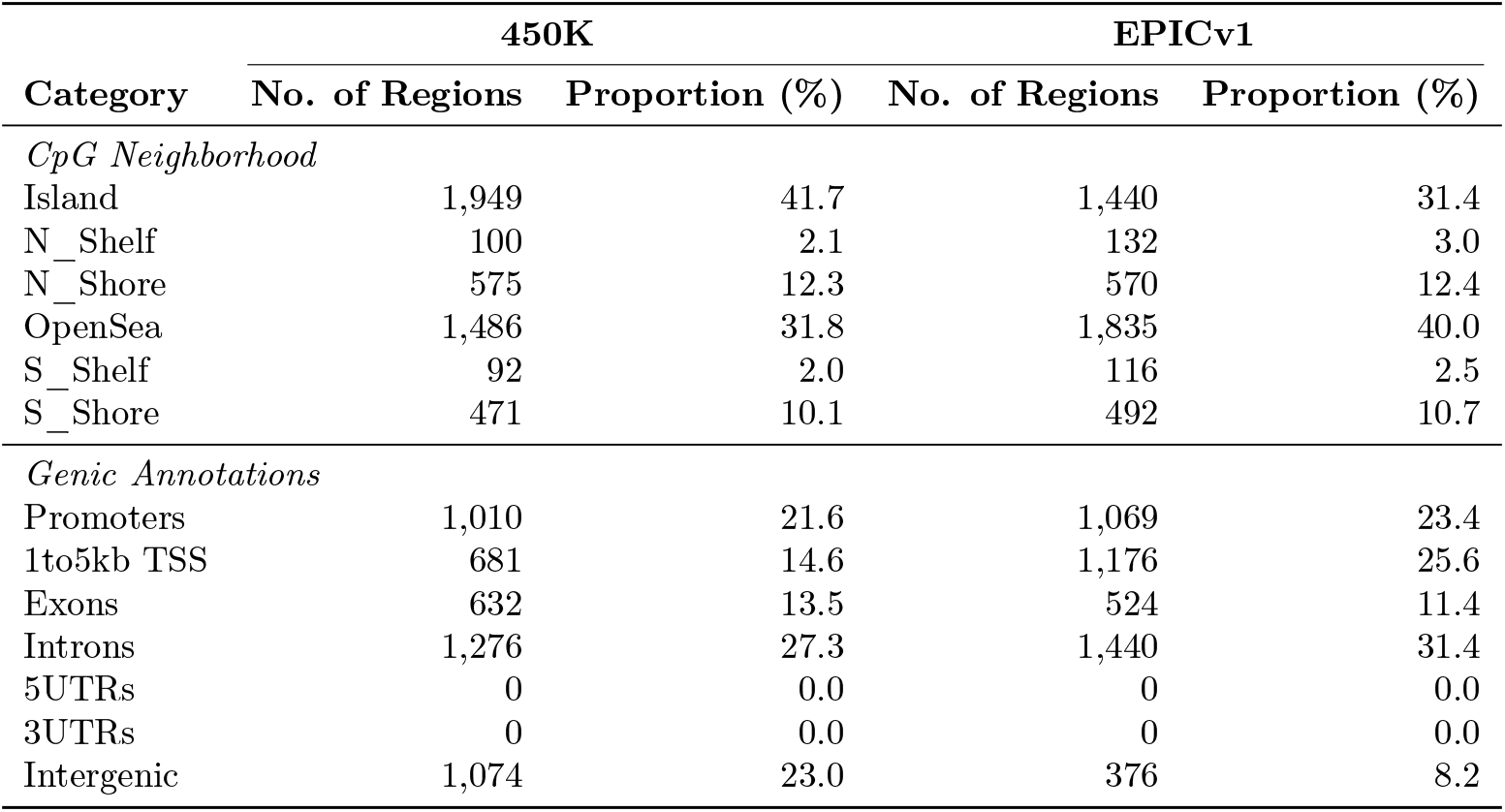
The table summarizes the distribution of co-methylated regions (*ρ* > 0.4) across CpG neighborhood classes and genic annotations for both 450K and EPICv1 arrays. The number of regions indicates the count of co-methylated regions in each genomic category and the proportion of regions represents the percentage of these regions relative to the total number of co-methylated regions identified on that array. CpG neighborhoods include Islands, Shores, Shelves, and OpenSea regions and genic annotations comprise of promoters, exons, introns and intergenic regions.

| Category | 450K |  | EPICv1 |  |
| --- | --- | --- | --- | --- |
|  | No. of Regions | Proportion (%) | No. of Regions | Proportion (%) |
| *CpG Neighborhood* |  |  |  |  |
| Island | 1,949 | 41.7 | 1,440 | 31.4 |
| N_Shelf | 100 | 2.1 | 132 | 3.0 |
| N_Shore | 575 | 12.3 | 570 | 12.4 |
| OpenSea | 1,486 | 31.8 | 1,835 | 40.0 |
| S_Shelf | 92 | 2.0 | 116 | 2.5 |
| S_Shore | 471 | 10.1 | 492 | 10.7 |
| *Genic Annotations* |  |  |  |  |
| Promoters | 1,010 | 21.6 | 1,069 | 23.4 |
| 1to5kb TSS | 681 | 14.6 | 1,176 | 25.6 |
| Exons | 632 | 13.5 | 524 | 11.4 |
| Introns | 1,276 | 27.3 | 1,440 | 31.4 |
| 5UTRs | 0 | 0.0 | 0 | 0.0 |
| 3UTRs | 0 | 0.0 | 0 | 0.0 |
| Intergenic | 1,074 | 23.0 | 376 | 8.2 |

### 3.2 SACOMA achieves optimal balance between sensitivity and specificity in detecting co-methlyation regions compared to popular unsupervised models

We evaluated SACOMA alongside existing models such as Aclust2 [17] and coMethDMR [18] to benchmark their ability to accurately identify co-methylated regions in DNAm array data. To comprehensively assess performance, we examined type-I error, true positives (TPs), false positives (FPs), false negatives (FNs) detection rates, false discovery rates (FDR) and statistical power across multiple simulation conditions that varied by array platform (450K and EPICv1), maximum inter-CpG gap (200 bp, 500 bp, and 1000 bp) and number of CpGs per region (3, 5, and 7). Under null simulations where no truly co-methylated regions exist any detected signal represents a false positive, making type-I error analysis crucial for evaluating model validity. Across all tested configurations, SACOMA along with Aclust2 consistently maintained strict type-I error control, remaining well below the 5% nominal threshold in both 450K and EPICv1 datasets, regardless of CpG density or gap size (**STable 3**). In contrast, coMethDMR displayed inflated false positive rates as the number of CpGs per region increased, with type-I error surpassing 10% in some 450K scenarios (0.105 at 200 bp and 0.139 at 500 bp). Even when the maximum inter-CpG gap was expanded to 1000 bp, SACOMA remained well-calibrated, while coMethDMR continued to exhibit instability (**STable 4**). These findings demonstrate SACOMA’s ability to control spurious detections and ensure robust inference, a key requirement for biologically interpretable co-methylation mapping.

Moving beyond type-I error control, we next assessed each model’s performance in detecting true co-methylated regions. For regions containing three CpGs, SACOMA not only matched Aclust2 in maintaining FPs but also detected substantially higher TPs and fewer FNs **(Figure 2A–D)**. For example, SACOMA detected 1,662 TPs under the 450K–200 bp configuration and 2,097 under EPICv1–200 bp, while maintaining FPs. In contrast, coMethDMR detected more TPs overall but at the cost of excessive FPs (437 under EPICv1–200 bp and 478 under EPICv1–500 bp), reducing its specificity. This pattern suggests that coMethDMR, by identifying a large number of regions without adequate false-positive control, effectively performs little to no dimension reduction capturing spurious, noise-driven regions alongside true signals.

**Figure 2.**
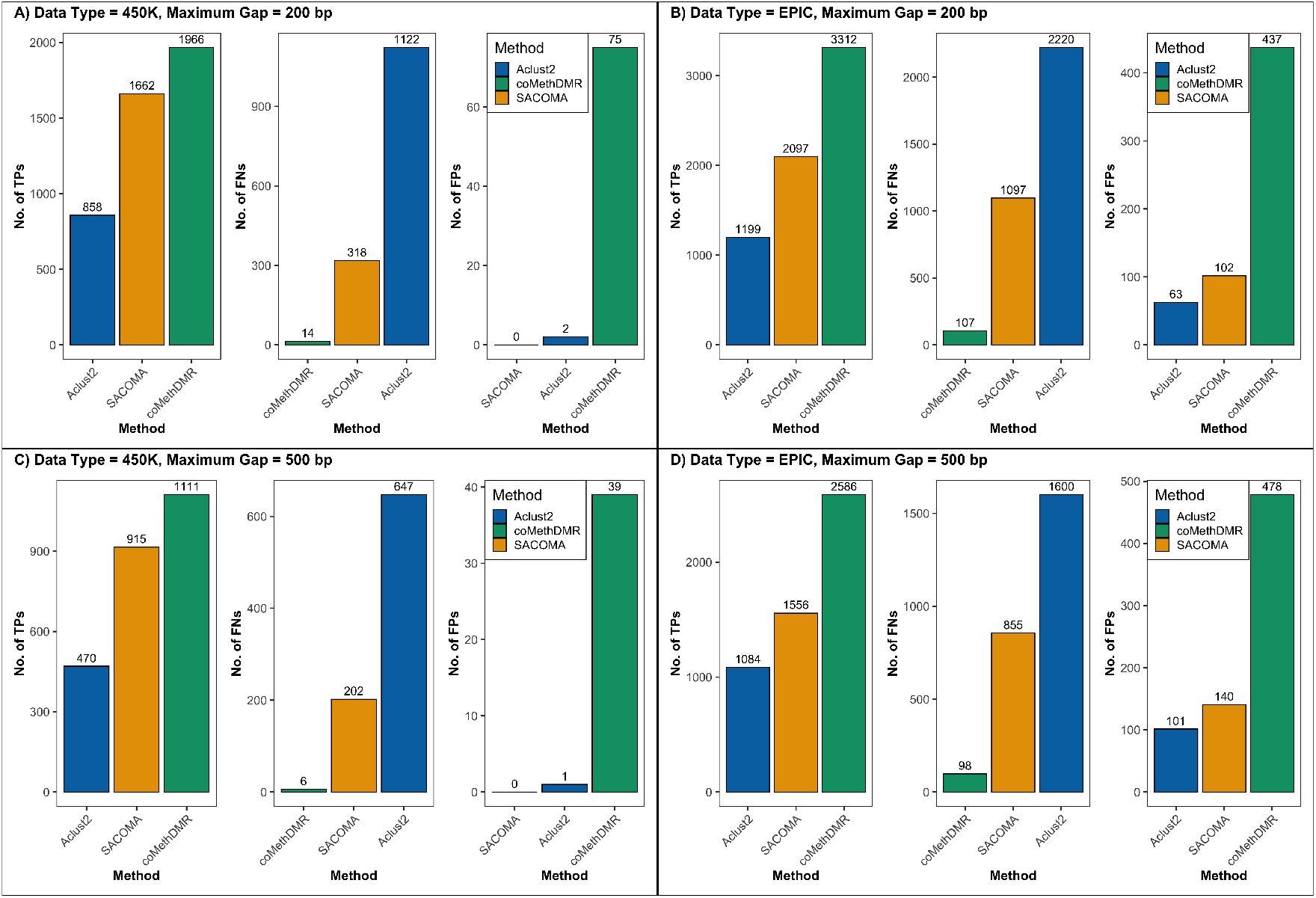
Performance evaluation of co-methylation models with 3 CpGs per region. This panel plot illustrates the number of True Positives (TPs), True Negatives (TNs) and False Positives (FPs) identified by three methods (Aclust2, SACOMA and coMethDMR) under different simulation settings. Results are shown for both the 450K and EPICv1 array templates with maximum gaps of 200 bp and 500 bp between CpGs. Panel (**A**) shows results for the 450K dataset with maxGap = 200 bp, panel (**B**) for EPICv1 with maxGap = 200 bp, panel (**C**) for 450K with maxGap = 500 bp, and panel (**D**) for EPICv1 with maxGap = 500 bp. These results are based on the median of 100 simulations. Models achieving higher numbers of TPs while maintaining low FPs demonstrate superior balance between sensitivity and specificity.

Such behavior not only increases the multiple-testing burden but can also propagate false discoveries in downstream region-based analyses. On the other hand, Aclust2, while maintaining low FP rates, produced comparatively fewer TPs, indicating that it may over-reduce by defining overly restrictive regions. This conservative behavior can lead to information loss and consequently low power in subsequent association analyses. As the number of CpGs per region increased to five and seven (**SFigure 1–2**), SACOMA continued to maintain low FP counts while preserving strong sensitivity, effectively balancing TPs and FNs. Aclust2 showed reduced sensitivity, whereas coMethDMR again demonstrated high FP counts across both array types. This robust pattern persisted under the expanded 1000 bp maximum gap condition (**SFigure 3**), confirming the stability of SACOMA’s performance across genomic contexts. In terms of FDR and statistical power, SACOMA achieved on average 40–50% higher power than Aclust2 across both platforms and all maximum gap settings (**STable 5** and **STable 6**), while keeping the FDR below 5% in most cases. In contrast, coMethDMR exhibited nearly perfect power (≥ 0.98 across most configurations), but this came at the cost of inflated FDR values (ranging from 0.09 to 0.18). This pattern suggests that coMethDMR, while identifying a large number of regions, performs little to no dimension reduction and captures spurious, noise-driven regions alongside true signals. Such behavior not only increases the multiple-testing burden but also undermines downstream region-based analyses by propagating false discoveries. Conversely, Aclust2, despite maintaining FDR control in most cases, displayed the lowest power across all settings (0.15–0.49), suggesting over-reduction defining overly restrictive regions that exclude many true co-methylated CpGs and lead to lower statistical power. In contrast, SACOMA demonstrated a balanced and robust performance, achieving effective dimension reduction while accurately identifying true co-methylated regions. This balance between power, FDR control and biological plausibility highlights SACOMA’s strength as a statistically rigorous and biologically coherent framework for identifying co-methylated regions across genome-wide datasets.

### 3.3 Real Data Analysis

In addition to simulation studies, we next applied SACOMA to two population-level DNAm datasets: 450K and EPICv1 (for more details, see Methods). For the 450K dataset **(Table 2)**, SACOMA identified 2,595 regions, with the number of CpGs per region ranging from 3 to 82, using a maximum gap of 200 bp between CpGs. In comparison, Aclust2 identified 1,736 regions and coMethDMR identified 4,551 regions. Importantly, 854 regions were consistently detected by all three models, underscoring the strong consistency across their outputs. Among the total co-methylated regions identified by SACOMA, 66% overlapped with coMethDMR and 39% overlapped with Aclust2, indicating that SACOMA captures biologically coherent regions that align well with other established unsupervised methods. Beyond this shared subset, SACOMA uniquely identified 712 regions, Aclust2 identified 690 and coMethDMR detected 2,810 regions. The disproportionately high number of unique regions called by coMethDMR suggests a strong likelihood of false positives, while SACOMA’s balanced detection indicates better alignment with biological plausibility. From a computational perspective, Aclust2 was the fastest to run, SACOMA maintained moderate runtime and coMethDMR was the most computationally intensive. Overall, coMethDMR’s behavior across datasets, including its lack of false-positive control, limited dimension reduction, high computational cost and strong tendency to identify an excessive number of regions, raises concerns about its overall reliability. In contrast, SACOMA demonstrated consistent, stable and computationally efficient performance across datasets.

We also investigated the genomic distribution of the regions identified by each model in the 450K dataset. SACOMA and Aclust2 showed broadly similar annotation patterns, with the majority of regions mapping to CpG islands (40.8% and 41.5%, respectively) and open-sea regions (25.6% and 26.9%). These distributions closely mirror the enrichment patterns observed in Section 3.1, where co-methylated regions were shown to predominantly occur within CpG islands and promoters. This correspondence reinforces that SACOMA effectively captures biologically informed co-methylation structure, consistent with genome-wide methylation organization. Compared to these models, coMethDMR displayed a slightly higher proportion of island-associated regions (44.2%), a pattern that aligns with its overall tendency to report a greater number of regions. SACOMA also demonstrated strong biological relevance in genic annotations (**SFigure 4**), with approximately one-third of its regions mapping to introns (32.8%) and another one-third to promoters (35.8%), aligning well with expected functional enrichment patterns. Together, these real-data results highlight the capability of SACOMA to identify co-methylated regions that are both statistically robust and biologically meaningful. Its ability to generalize across array platforms while preserving annotation concordance underscores its practical utility for epigenomic studies. The results of the EPICv1 dataset were consistent with these findings and are provided in the Supplementary Materials **STable 7** and **SFigure 5**.

**Table 2.** The table presents a comparison of performance metrics, genomic region characteristics, and computational resource usage across three models (SACOMA, Aclust2 and coMethDMR) applied to the 450K dataset. The table summarizes the total number of regions detected, CpGs per region, number of unique and common regions, peak memory consumption, and total runtime. Additionally, the lower section presents the genomic context distribution of the identified regions (Island, Shore, Shelf, and OpenSea) with both counts and percentages. Counts represent the number of regions in each CpG neighborhood and the percentages (in parentheses) are their proportions out of the total number of regions identified by each model. Memory usage is reported in gigabytes (GB) and time in minutes.

| Metric | SACOMA | Aclust2 | coMethDMR |
| --- | --- | --- | --- |
| No. of Regions | 2595 | 1736 | 4551 |
| No. of CpGs/Region | [3, 82] | [3, 45] | [3, 82] |
| No. of Unique Regions | 712 | 690 | 2810 |
| Common Regions | 854 | 854 | 854 |
| Peak Memory Usage (GB) | 0.893 | 1.675 | 0.652 |
| Runtime (min) | 76.87 | 4.99 | 128.58 |
| *CpG Neighborhood* |  |  |  |
| Island | 1060 (40.8%) | 721 (41.5%) | 2011 (44.2%) |
| N_Shelf | 56 (2.2%) | 35 (2%) | 109 (2.4%) |
| N_Shore | 405 (15.6%) | 257 (14.8%) | 649 (14.3%) |
| OpenSea | 665 (25.6%) | 467 (26.9%) | 1162 (25.5%) |
| S_Shelf | 45 (1.8%) | 30 (1.8%) | 84 (1.8%) |
| S_Shore | 364 (14%) | 226 (13%) | 536 (11.8%) |

## 4 Discussion

We introduce the SACOMA algorithm which is based on spatially constrained hierarchical clustering designed to group adjacent DNAm sites according to both their spatial proximity and similarity in methylation patterns. The method formulates region detection in DNA methyation data as a hierarchical clustering problem that jointly minimizes differences within clusters in terms of methylation levels and genomic distance. Because co-methylated regions are typically defined by local correlation and spatial closeness yet vary in size and structure, SACOMA integrates these two factors through a tunable mixing parameter. This allows the algorithm to identify co-methylated regions in a flexible, data-driven framework without the dependence on rigid assumptions and sensitivity to hyperparameter specifications. SACOMA serves two main purposes: (1) detecting regions of co-regulated methylation sites, and (2) reducing data dimensionality by aggregating sites into a smaller set of meaningful analytical units that can be used for downstream region-based association analysis such as in EWAS. Comprehensive simulation studies demonstrated that SACOMA outperforms existing unsupervised methods in sensitivity and specificity for accurately identifying co-methylated regions using Illumina Infinium 450K and 850K BeadChip data. Furthermore, applying SACOMA to population-level methylation datasets revealed its ability to identify co-methylated regions with significant functional and biological relevance.

In simulation-based comparison with existing unsupervised methods, SACOMA consistently achieved high sensitivity while maintaining strict control of type-I error and false positive rates across varying CpG densities and genomic gap thresholds. In contrast, coMethDMR, although capable of detecting a large number of regions, frequently exhibited inflated false positives suggesting that the higher counts of detected regions may not always represent true co-methylated domains but rather noise or spurious associations. This reinforces that a greater number of identified regions does not necessarily equate to higher biological accuracy. Moreover, the process of identifying co-methylated regions can be viewed as a form of dimension reduction, condensing thousands of CpG-level signals into a smaller set of interpretable regional features. While this approach enhances computational efficiency, models that over-reduce may lose biologically relevant signals, whereas those that under-reduce may introduce noise and redundancy. SACOMA achieves an optimal balance by adaptively learning CpG clusters that are both compact and homogeneous, thus providing an efficient yet proper representation of methylation structure.

In the analysis of two population-level DNAm array datasets from two different platforms (450K and EPICv1), SACOMA demonstrated robust, biologically coherent performance. It identified regions enriched in CpG islands and promoter-associated domains, genomic contexts well known for coordinated methylation regulation and functional importance in gene expression [38, 14, 2, 4]. Approximately one-third of SACOMA-identified regions mapped to promoters and another third to introns, reflecting patterns consistent with established genomic organization of methylation. These results mirror earlier findings that CpG islands are frequent hotspots for co-methylation and play critical roles in transcriptional regulation and chromatin state maintenance [39, 4]. Compared to other models, SACOMA showed stronger biological plausibility, moderate computational cost, and reproducible performance across both array platforms, reaffirming its practical utility in real-world methylation studies.

Although SACOMA has been developed within the context of DNAm array studies, its underlying framework is broadly applicable to a wide range of data types that display spatial or structured dependence. Because SACOMA models correlations among spatially ordered features while optimizing local homogeneity, it can also be adapted for sequencing-based DNAm data such as whole-genome or reduced representation bisulfite sequencing, where methylation levels exhibit spatial continuity along the genome. Given the denser CpG coverage in these sequencing platforms compared to the 450K or EPIC arrays, SACOMA may perform even more effectively by leveraging richer correlation structure and improved spatial resolution. Beyond methylation, SACOMA can be extended to other epigenomic and multi-omic datasets with similar architectures, including chromatin accessibility data from ATAC-seq, where genomic loci show correlated accessibility patterns [40], and histone modification profiles from ChIP-seq, which often display spatially coherent enrichment domains [41]. The framework can further be applied outside the field of omics to structured clinical datasets such as electronic health records, where features may show spatial or temporal correlation, for example through geographic clustering or longitudinal dependencies across repeated measurements [42]. In addition, SACOMA can be coupled with any downstream association or predictive modeling approach, as the co-methylated regions it identifies serve as stable and biologically informed units of analysis that enhance both statistical power and interpretability. Future studies may expand on this framework by integrating SACOMA with more sophisticated modeling strategies to further improve detection sensitivity and capture complex spatial or correlation structures.

While SACOMA provides a strong and adaptable framework for identifying co-methylated regions, some limitations should be recognized. Firstly, SACOMA relies on hierarchical spatial clustering, which can be sensitive to differences in local CpG density and variation in correlation strength. In regions where CpG coverage is limited, such as in Illumina 450K and EPIC arrays, this dependence may lead to fragmented or unstable clusters. This issue can be mitigated by employing adaptive distance thresholds that stabilize clustering in sparse genomic contexts. Secondly, hierarchical clustering methods have difficulty handling missing values [43]. This challenge can be addressed through imputation approaches specifically designed for DNAm data, such as methyLImp or missForest [44], which leverage local correlation structures and methylation patterns to accurately recover missing beta values [45]. Finally, when applied to genome-wide methylation data, which are inherently high dimensional, clustering algorithms may lose sensitivity because the contrast between near and far distances diminishes with increasing dimensionality, making global distance-based clustering less meaningful [43]. SACOMA addresses this challenge by first partitioning the genome into smaller, spatially constrained CpG bins based on local proximity. This localized approach ensures that distance measures remain informative within each bin, effectively reducing dimensionality and computational complexity while preserving biologically relevant methylation patterns.

In conclusion, SACOMA addresses a crucial methodological gap in unsupervised region-based DNAm array analysis by offering a biologically informed and statistically robust framework for identifying co-methylated regions, serving as a foundational step for region-level analysis through an unsupervised framework. Through an adaptive co-methylated region detection, it achieves an effective balance between sensitivity, specificity, and computational efficiency. By improving this foundational analysis stage, SACOMA enables more accurate, interpretable, and biologically meaningful downstream analysis which holds the potential to provide insights into epigenetic regulation, establishing itself as a powerful and generalizable tool for future large-scale methylation studies.

## Availability of Data and Materials

The 450K dataset is accessible from the GEO database under accession number GSE281199. The EPICv1 dataset is accessible from the GEO database under accession number GSE169338. Code for SACOMA, simulations, and comparisons of methods identifying co-methylated regions on the various datasets explored are available from GitHub at https://github.com/SChatLab/SACOMA. The source code is licensed under an MIT License.

## Author Contributions

S.C. conceived and designed this study; S.M. gathered and managed data; S.M. analyzed the data; S.C., S.M. and A.F. wrote the draft paper and interpreted the results; S.C. supervised the work. A.G. provided critical intellectual content. All authors approved the final manuscript.

## Acknowledgments

This research was supported in part by Lilly Endowment, Inc., through its support for the Indiana University Pervasive Technology Institute.

## Competing Interests

The authors declare that they have no competing interests.

## Notes

### Competing Interest Statement

The authors have declared no competing interest.

## References

[1] Rudolf Jaenisch and Adrian Bird. Epigenetic regulation of gene expression: how the genome integrates intrinsic and environmental signals. Nature genetics, 33(3):245–254, 2003.

[2] Jordana T Bell, Athma A Pai, Joseph K Pickrell, Daniel J Gaffney, Roger Pique-Regi, Jacob F Degner, Yoav Gilad, and Jonathan K Pritchard. Dna methylation patterns associate with genetic and gene expression variation in hapmap cell lines. Genome biology, 12(1):R10, 2011.

[3] Manel Esteller. Epigenetics in cancer. New England Journal of Medicine, 358(11):1148–1159, 2008.

[4] Aimée M Deaton and Adrian Bird. Cpg islands and the regulation of transcription. Genes & development, 25(10):1010–1022, 2011.

[5] Vardhman K Rakyan, Thomas A Down, David J Balding, and Stephan Beck. Epigenome-wide association studies for common human diseases. Nature Reviews Genetics, 12(8):529–541, 2011.

[6] Christine Ladd-Acosta. Epigenetic signatures as biomarkers of exposure. Current environmental health reports, 2(2):117–125, 2015.

[7] Bonnie R Joubert, Janine F Felix, Paul Yousefi, Kelly M Bakulski, Allan C Just, Carrie Breton, Sarah E Reese, Christina A Markunas, Rebecca C Richmond, Cheng-Jian Xu, et al. Dna methylation in newborns and maternal smoking in pregnancy: genome-wide consortium meta-analysis. The American Journal of Human Genetics, 98(4):680–696, 2016.

[8] Marina Bibikova, Bret Barnes, Chan Tsan, Vincent Ho, Brandy Klotzle, Jennie M Le, David Delano, Lu Zhang, Gary P Schroth, Kevin L Gunderson, et al. High density dna methylation array with single cpg site resolution. Genomics, 98(4):288–295, 2011.

[9] Ruth Pidsley, Elena Zotenko, Timothy J Peters, Mitchell G Lawrence, Gail P Risbridger, Peter Molloy, Susan Van Djik, Beverly Muhlhausler, Clare Stirzaker, and Susan J Clark. Critical evaluation of the illumina methylationepic beadchip microarray for whole-genome dna methylation profiling. Genome biology, 17(1):208, 2016.

[10] James M Flanagan. Epigenome-wide association studies (ewas): past, present, and future. Cancer Epigenetics: Risk Assessment, Diagnosis, Treatment, and Prognosis, pages 51–63, 2014.

[11] Yun Liu, Martin J Aryee, Leonid Padyukov, M Daniele Fallin, Espen Hesselberg, Arni Runarsson, Lovisa Reinius, Nathalie Acevedo, Margaret Taub, Marcus Ronninger, et al. Epigenome-wide association data implicate dna methylation as an intermediary of genetic risk in rheumatoid arthritis. Nature biotechnology, 31(2):142–147, 2013.

[12] Timothy J Peters, Michael J Buckley, Aaron L Statham, Ruth Pidsley, Katherine Samaras, Reginald V Lord, Susan J Clark, and Peter L Molloy. De novo identification of differentially methylated regions in the human genome. Epigenetics & chromatin, 8(1):6, 2015.

[13] Andrew E Jaffe, Peter Murakami, Hwajin Lee, Jeffrey T Leek, M Daniele Fallin, Andrew P Feinberg, and Rafael A Irizarry. Bump hunting to identify differentially methylated regions in epigenetic epidemiology studies. International journal of epidemiology, 41(1):200–209, 2012.

[14] Ornella Affinito, Domenico Palumbo, Annalisa Fierro, Mariella Cuomo, Giulia De Riso, Antonella Monticelli, Gennaro Miele, Lorenzo Chiariotti, and Sergio Cocozza. Nucleotide distance influences co-methylation between nearby cpg sites. Genomics, 112(1):144–150, 2020.

[15] Florian Eckhardt, Joern Lewin, Rene Cortese, Vardhman K Rakyan, John Attwood, Matthias Burger, John Burton, Tony V Cox, Rob Davies, Thomas A Down, et al. Dna methylation profiling of human chromosomes 6, 20 and 22. Nature genetics, 38(12):1378–1385, 2006.

[16] Tamar Sofer, Elizabeth D Schifano, Jane A Hoppin, Lifang Hou, and Andrea A Baccarelli. A-clustering: a novel method for the detection of co-regulated methylation regions, and regions associated with exposure. Bioinformatics, 29(22):2884–2891, 2013.

[17] Oladele A Oluwayiose, Haotian Wu, Feng Gao, Andrea A Baccarelli, Tamar Sofer, and J Richard Pilsner. Aclust2. 0: a revamped unsupervised r tool for infinium methylation beadchips data analyses. Bioinformatics, 38(20):4820–4822, 2022.

[18] Lissette Gomez, Gabriel J Odom, Juan I Young, Eden R Martin, Lizhong Liu, Xi Chen, Anthony J Griswold, Zhen Gao, Lanyu Zhang, and Lily Wang. comethdmr: accurate identification of co-methylated and differentially methylated regions in epigenome-wide association studies with continuous phenotypes. Nucleic acids research, 47(17):e98–e98, 2019.

[19] Luc Anselin. Local indicators of spatial association—lisa. Geographical analysis, 27(2):93–115, 1995.

[20] Martin Ester, Hans-Peter Kriegel, Jörg Sander, and Xiaowei Xu. A density-based algorithm for discovering clusters in large spatial databases with noise. In Proceedings of the Second International Conference on Knowledge Discovery and Data Mining, pages 226–231. AAAI Press, 1996.

[21] Derya Birant and Alp Kut. St-dbscan: An algorithm for clustering spatial–temporal data. Data & Knowledge Engineering, 60(1):208–221, 2007.

[22] Juan C Duque, Raul Ramos, and Jordi Suriñach. Supervised regionalization methods: A survey. International Regional Science Review, 35(4):451–479, 2012.

[23] Fernando Bacao, Victor Lobo, and Marco Painho. Self-organizing maps as substitutes for k-means clustering. Computers & Geosciences, 31(5):531–544, 2005.

[24] Marie Chavent, Vanessa Kuentz-Simonet, Amaury Labenne, and Jérôme Saracco. Clustgeo: an r package for hierarchical clustering with spatial constraints. Computational Statistics, 33(4):1799–1823, 2018.

[25] Rafael G. Cavalcante and M. A. Sartor. annotatr: Genomic regions in context, 2017. R package version 1.34.0.

[26] Michael Lawrence, Wolfgang Huber, Hervé Pagès, Patrick Aboyoun, Marc Carlson, Robert Gentleman, Martin T Morgan, and Vincent J Carey. Software for computing and annotating genomic ranges. PLoS computational biology, 9(8):e1003118, 2013.

[27] Bioconductor Package Maintainer. TxDb.Hsapiens.UCSC.hg19.knownGene: Annotation package for TxDb object(s), 2025. R package version 3.2.2.

[28] Bioconductor Package Maintainer. TxDb.Hsapiens.UCSC.hg38.knownGene: Annotation package for TxDb object(s), 2025. R package version 3.18.0.

[29] Zheng (Joe) Dong, Samantha Schaffner, Maggie Fu, Joanne Whitehead, Julia L. MacIsaac, David H. Rehkopf, W. Thomas Boyce, Luis Rosero-Bixby, Lluis Quintana-Murci, Etienne Patin, Gregory E. Miller, Keegan Korthauer, and Michael S. Kobor. Genetic basis, quantitative nature, and functional relevance of evolutionarily conserved dna methylation. bioRxiv, 2024.

[30] Martin J Aryee, Andrew E Jaffe, Hector Corrada-Bravo, Christine Ladd-Acosta, Andrew P Feinberg, Kasper D Hansen, and Rafael A Irizarry. Minfi: a flexible and comprehensive bioconductor package for the analysis of infinium dna methylation microarrays. Bioinformatics, 30(10):1363–1369, 2014.

[31] Sean Davis and Sven Bilke. An introduction to the methylumi package. Biocond. Package, 10:B9, 2010.

[32] Andrew E Teschendorff, Francesco Marabita, Matthias Lechner, Thomas Bartlett, Jesper Tegner, David Gomez-Cabrero, and Stephan Beck. A beta-mixture quantile normalization method for correcting probe design bias in illumina infinium 450 k dna methylation data. Bioinformatics, 29(2):189–196, 2013.

[33] W Evan Johnson, Cheng Li, and Ariel Rabinovic. Adjusting batch effects in microarray expression data using empirical bayes methods. Biostatistics, 8(1):118–127, 2007.

[34] Jeffrey T. Leek, W. Evan Johnson, Hilary S. Parker, Elana J. Fertig, Andrew E. Jaffe, Yuqing Zhang, John D. Storey, and Leonardo Collado Torres. sva: Surrogate Variable Analysis, 2025. R package version 3.56.0.

[35] Jagyashila Das, Nitya Wadhwa, Uma CM Natchu, Ramachandran Thiruvengadam, Pallavi Kshetrapal, Shinjini Bhatnagar, Partha P Majumder, and Arindam Maitra. Genome-wide temporal landscaping of dna methylation in pregnant women delivering at term: a garbh-ini study. Epigenomics, 15(9):543–556, 2023.

[36] Tiffany J Morris, Lee M Butcher, Andrew Feber, Andrew E Teschendorff, Ankur R Chakravarthy, Tomasz K Wojdacz, and Stephan Beck. Champ: 450k chip analysis methylation pipeline. Bioinformatics, 30(3):428–430, 2014.

[37] Charles Spearman. The proof and measurement of association between two things. 1961.

[38] Olga Taryma-Leśniak, Jan Bińkowski, Patrycja Kamila Przybylowicz, Katarzyna Ewa Sokolowska, Konrad Borowski, and Tomasz Kazimierz Wojdacz. Methylation patterns at the adjacent cpg sites within enhancers are a part of cell identity. Epigenetics & Chromatin, 17(1):30, 2024.

[39] Tapas K Kundu. Cpg islands in chromatin organization and gene expression. The Journal of Biochemistry, 125(2):217–222, 1999.

[40] Rachel E Gate, Christine S Cheng, Aviva P Aiden, Atsede Siba, Marcin Tabaka, Dmytro Lituiev, Ido Machol, M Grace Gordon, Meena Subramaniam, Muhammad Shamim, et al. Genetic determinants of co-accessible chromatin regions in activated t cells across humans. Nature genetics, 50(8):1140–1150, 2018.

[41] Shiliyang Xu, Sean Grullon, Kai Ge, and Weiqun Peng. Spatial clustering for identification of chip-enriched regions (sicer) to map regions of histone methylation patterns in embryonic stem cells. In Stem cell transcriptional networks: methods and protocols, pages 97–111. Springer, 2014.

[42] Abolfazl Mollalo, Bashir Hamidi, Leslie A Lenert, and Alexander V Alekseyenko. Application of spatial analysis on electronic health records to characterize patient phenotypes: systematic review. JMIR Medical Informatics, 12(1):e56343, 2024.

[43] Divya Pandove, Shivan Goel, and Rinkl Rani. Systematic review of clustering high-dimensional and large datasets. ACM Transactions on Knowledge Discovery from Data (TKDD), 12(2):1–68, 2018.

[44] Daniel J Stekhoven and Peter Bühlmann. Missforest—non-parametric missing value imputation for mixed-type data. Bioinformatics, 28(1):112–118, 2012.

[45] Pietro Di Lena, Claudia Sala, Andrea Prodi, and Christine Nardini. Methylation data imputation performances under different representations and missingness patterns. BMC bioinformatics, 21(1):268, 2020.

