## Supplementary Materials for "A spatially-aware unsupervised pipeline to identify co-methylation regions in DNA methylation data"

#### Contents

**1 Supplementary Figures** **2**

**2 Supplementary Tables** **7**

### 1 Supplementary Figures

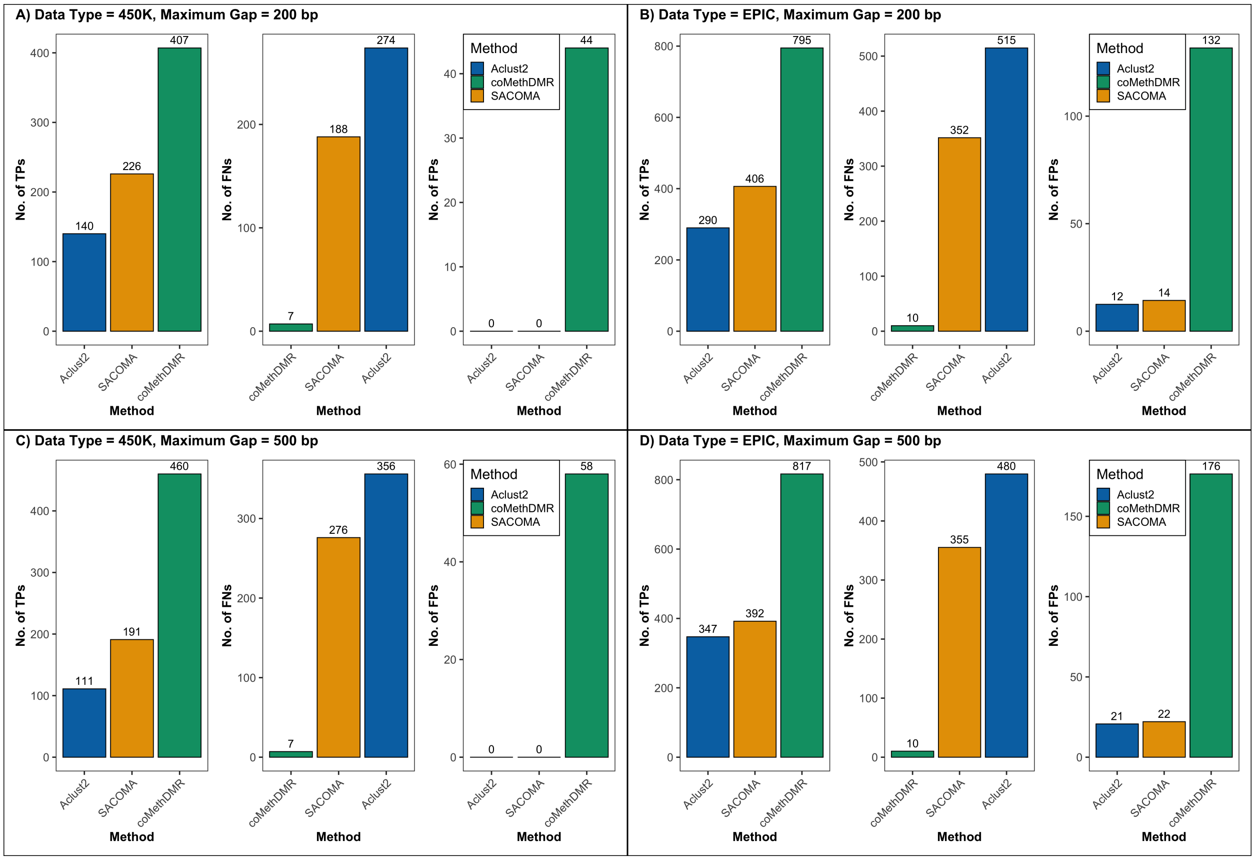


**SFigure 1: Performance evaluation of co-methylation models with 5 CpGs per region**. This panel plot illustrates the number of True Positives (TPs), True Negatives (TNs) and False Positives (FPs) identified by three methods (Aclust2, SACOMA and coMethDMR) under different simulation settings. Results are shown for both the 450K and EPICv1 array templates with maximum gaps of 200 bp and 500 bp between CpGs. Panel (**A**) shows results for the 450K dataset with maxGap = 200 bp, panel (**B**) for EPICv1 with maxGap = 200 bp, panel (**C**) for 450K with maxGap = 500 bp, and panel (**D**) for EPICv1 with maxGap = 500 bp. These results are based on the median of 100 simulations. Models achieving higher numbers of TPs while maintaining low FPs demonstrate superior balance between sensitivity and specificity.

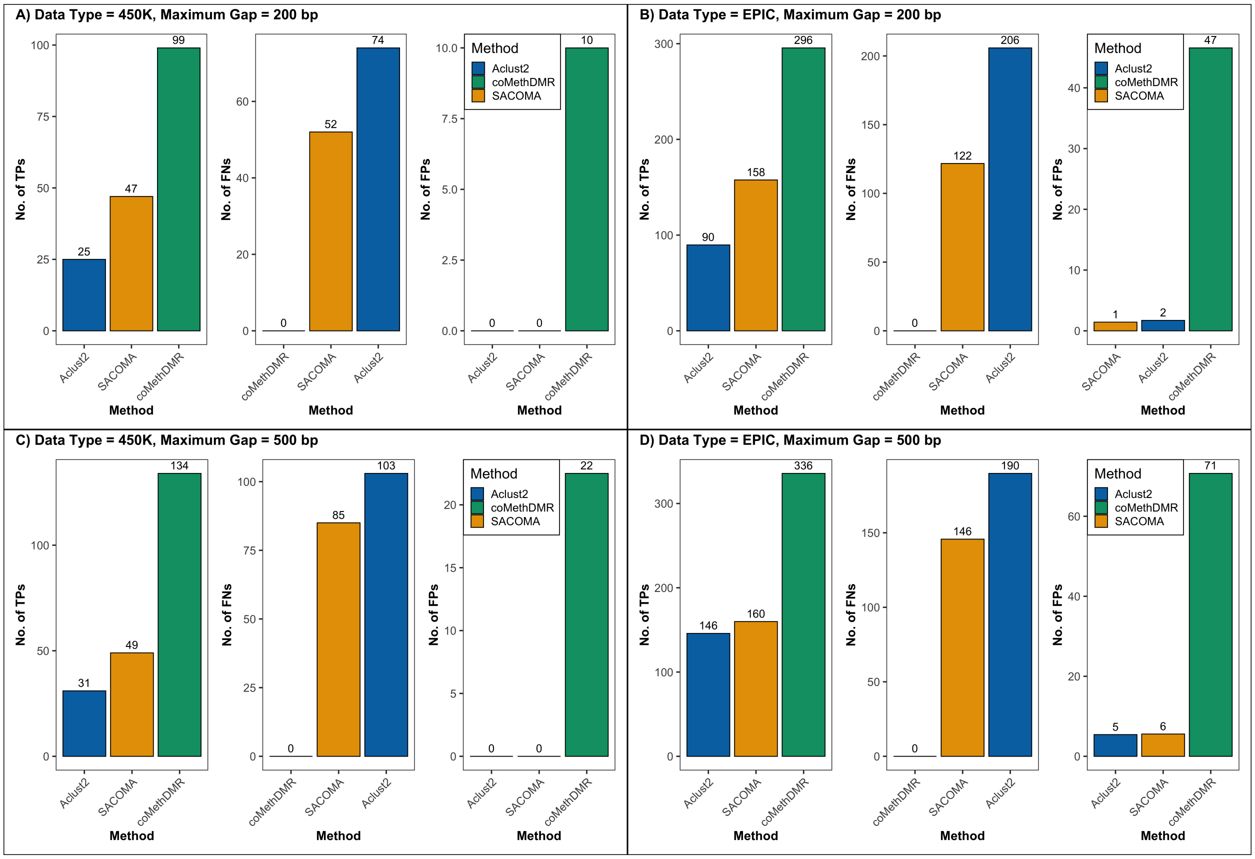


**SFigure 2: Performance evaluation of co-methylation models with 7 CpGs per region**. This panel plot illustrates the number of True Positives (TPs), True Negatives (TNs) and False Positives (FPs) identified by three methods (Aclust2, SACOMA and coMethDMR) under different simulation settings. Results are shown for both the 450K and EPICv1 array templates with maximum gaps of 200 bp and 500 bp between CpGs. Panel (**A**) shows results for the 450K dataset with maxGap = 200 bp, panel (**B**) for EPICv1 with maxGap = 200 bp, panel (**C**) for 450K with maxGap = 500 bp, and panel (**D**) for EPICv1 with maxGap = 500 bp. These results are based on the median of 100 simulations. Models achieving higher numbers of TPs while maintaining low FPs demonstrate superior balance between sensitivity and specificity.

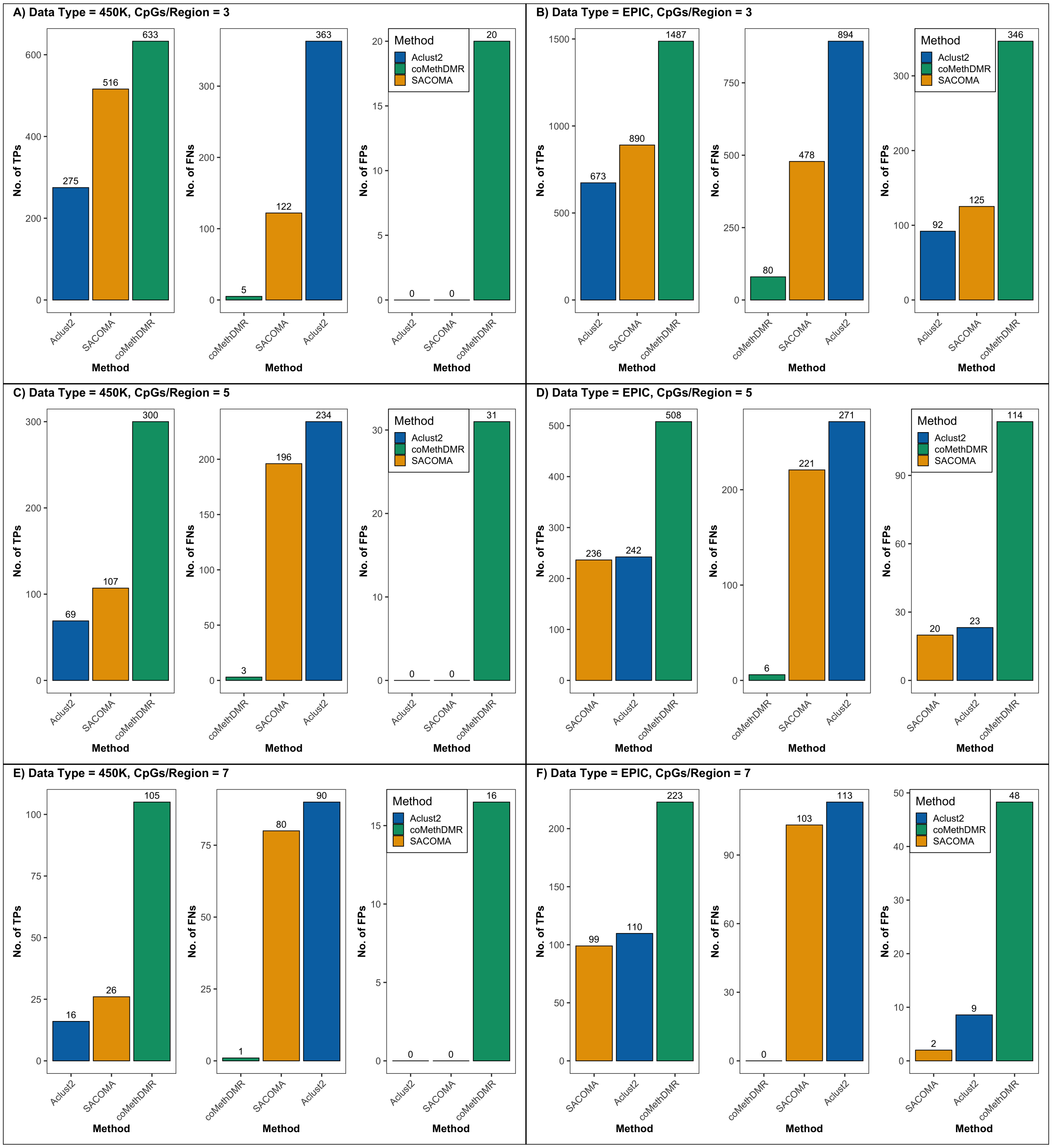


**SFigure 3: Performance evaluation of co-methylation models under a 1000 bp maximum gap condition**. This panel plot illustrates the number of True Positives (TPs), False Negatives (FNs), and False Positives (FPs) identified by three methods (Aclust2, SACOMA and coMethDMR) when applied to regions containing 3, 5, and 7 CpGs. Results are shown for both the 450K and EPICv1 array templates. Panels (**A**–**B**) display performance with 3 CpGs per region, (**C**–**D**) with 5 CpGs per region, and (**E**–**F**) with 7 CpGs per region. These results are based on the median of 100 simulations. Models achieving higher numbers of TPs while maintaining low FPs demonstrate superior balance between sensitivity and specificity.

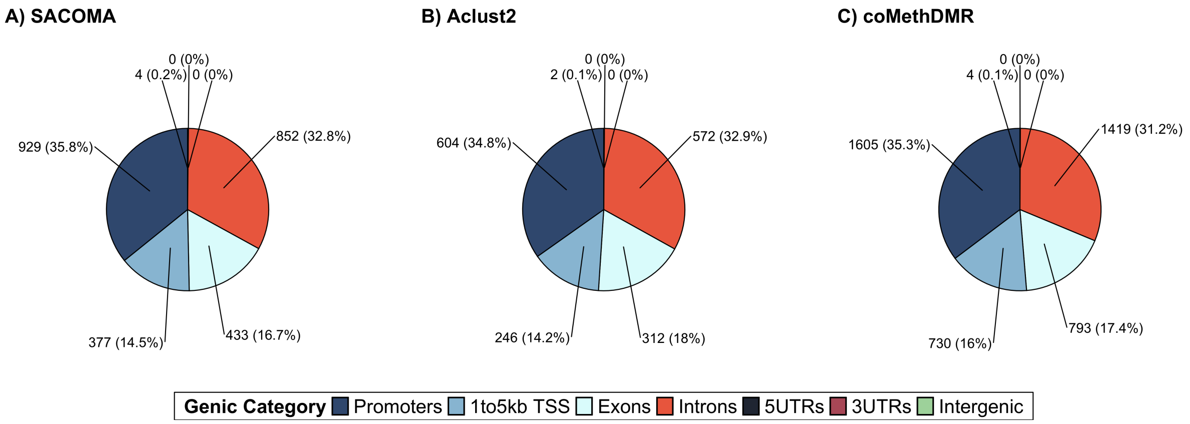


**SFigure 4: Genic annotation of identified regions in 450K real data analysis.** This panel plot shows the distribution of regions detected by four methods (SACOMA, Aclust2 and coMethDMR) across genic categories, such as promoters, TSS (1–5 kb), exons, introns, UTRs and intergenic regions. Panel (**A**) displays SACOMA, (**B**) Aclust2 and (**C**) coMethDMR.

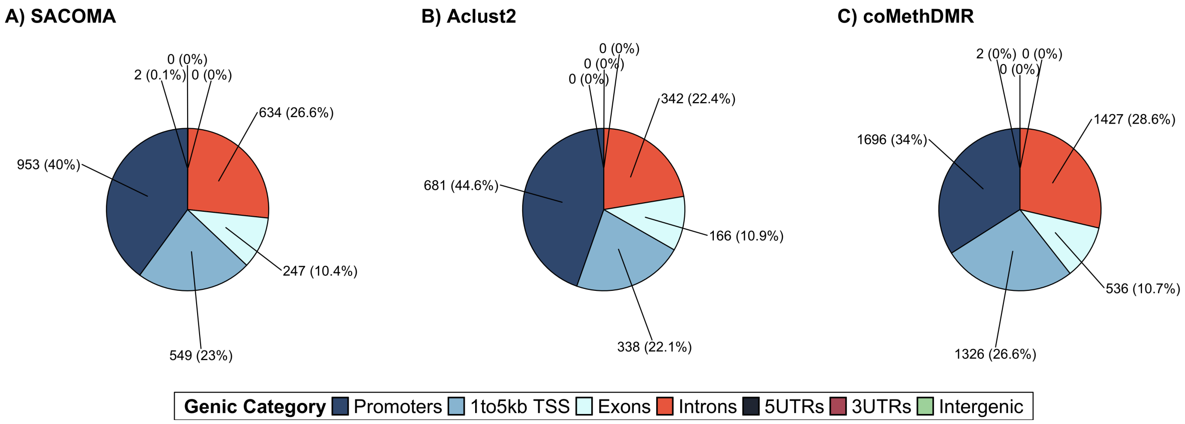


**SFigure 5: Genic annotation of identified regions in EPICv1 real data analysis.** This panel plot shows the distribution of regions detected by four methods (SACOMA, Aclust2 and coMethDMR) across genic categories, such as promoters, TSS (1–5 kb), exons, introns, UTRs, and intergenic regions. Panel (**A**) displays SACOMA, (**B**) Aclust2 and (**C**) coMethDMR

#### 2 Supplementary Tables

**STable 1: Summary of simulated regions under varying CpG densities and gap thresholds for 450K array template**. This table reports the total number of simulated regions, subdivided into co-methylated and non-co-methylated sets, for the 450K array template. Columns correspond to different maximum gap thresholds (200 bp, 500 bp and 1000 bp) and varying numbers of CpGs per region (3, 5 and 7). For each condition, the table provides the total regions simulated, along with an equal split between co-methylated and non-co-methylated regions. These settings define the benchmark scenarios used to evaluate model performance.

| | 450K | | | | | | | | |
|---|---|---|---|---|---|---|---|---|---|
| | maxGap = 200 | | | maxGap = 500 | | | maxGap = 1000 | | |
| | 3 | 5 | 7 | 3 | 5 | 7 | 3 | 5 | 7 |
| No. of Regions | 3960 | 828 | 198 | 2234 | 934 | 268 | 1276 | 606 | 212 |
| Co-Methylated Regions | 1980 | 414 | 99 | 1117 | 467 | 134 | 638 | 303 | 106 |
| Non-Co-Methylated Regions | 1980 | 414 | 99 | 1117 | 467 | 134 | 638 | 303 | 106 |

**STable 2: Summary of simulated regions under varying CpG densities and gap thresholds EPICv1 array template**. This table reports the total number of simulated regions, subdivided into co-methylated and non-co-methylated sets, for the EPICv1 array template. Columns correspond to different maximum gap thresholds (200 bp, 500 bp and 1000 bp) and varying numbers of CpGs per region (3, 5 and 7). For each condition, the table provides the total regions simulated, along with an equal split between co-methylated and non-co-methylated regions. These settings define the benchmark scenarios used to evaluate model performance.

| | EPICv1 | | | | | | | | |
|---|---|---|---|---|---|---|---|---|---|
| | maxGap = 200 | | | maxGap = 500 | | | maxGap = 1000 | | |
| | 3 | 5 | 7 | 3 | 5 | 7 | 3 | 5 | 7 |
| No. of Regions | 2814 | 518 | 220 | 1708 | 404 | 130 | 944 | 226 | 78 |
| Co-Methylated Regions | 1407 | 259 | 74 | 854 | 202 | 65 | 472 | 113 | 39 |
| Non-Co-Methylated Regions | 1407 | 259 | 74 | 854 | 202 | 65 | 472 | 113 | 39 |

**STable 3:** This table compares type-I error rates across three methods (Aclust2, SACOMA and coMethDMR) under different simulation settings. The first column lists the methods, while subsequent columns correspond to combinations of array type (450K and EPICv1), maximum gap between CpGs (200 bp and 500 bp) and number of CpGs per region (3, 5 and 7). Values represent the proportion of false positives detected under each condition. These results are based on the median of 100 simulations. Values exceeding the nominal 5% indicate inflated type-I error.

| Method | 450K | | | | | | EPICv1 | | | | | |
|---|---|---|---|---|---|---|---|---|---|---|---|---|
| | maxGap = 200 | | | maxGap = 500 | | | maxGap = 200 | | | maxGap = 500 | | |
| | 3 | 5 | 7 | 3 | 5 | 7 | 3 | 5 | 7 | 3 | 5 | 7 |
| Aclust2 | 0 | 0 | 0 | 0 | 0 | 0 | 0 | 0.001 | 0.002 | 0 | 0 | 0 |
| SACOMA | 0 | 0 | 0 | 0 | 0 | 0 | 0 | 0 | 0 | 0 | 0 | 0 |
| coMethDMR | 0.006 | 0.072 | 0.105 | 0.004 | 0.087 | 0.139 | 0.01 | 0.086 | 0.106 | 0.008 | 0.067 | 0.121 |

**STable 4:** This table compares type-I error rates across three methods (Aclust2, SACOMA and coMethDMR) under different simulation settings. The first column lists the methods, while subsequent columns correspond to combinations of array type (450K and EPICv1), maximum gap between CpGs (1000 bp) and number of CpGs per region (3, 5 or 7). Values represent the proportion of false positives detected under each condition. These results are based on the median of 100 simulations. Values exceeding the nominal 5% threshold indicate inflated type-I error.

| Method | 450K (maxGap = 1000) | | | EPICv1 (maxGap = 1000) | | |
|---|---|---|---|---|---|---|
| | 3 | 5 | 7 | 3 | 5 | 7 |
| Aclust2 | 0 | 0 | 0 | 0 | 0 | 0 |
| SACOMA | 0 | 0 | 0 | 0 | 0 | 0 |
| coMethDMR | 0.008 | 0.086 | 0.162 | 0.006 | 0.05 | 0.103 |

**STable 5:** Comparison of statistical power across models (Aclust2, SACOMA and coMethDMR) under varying minimum CpG thresholds (3, 5, 7) and maximum gap distances (200 bp, 500 bp, 1000 bp) for both Illumina 450K and EPICv1 arrays. These results are based on the median of 100 simulations.

| | **450K** | | | | | | | | | **EPICv1** | | | | | | | | |
|---|---|---|---|---|---|---|---|---|---|---|---|---|---|---|---|---|---|---|
| | maxGap = 200 | | | maxGap = 500 | | | maxGap = 1000 | | | maxGap = 200 | | | maxGap = 500 | | | maxGap = 1000 | | |
| | 3 | 5 | 7 | 3 | 5 | 7 | 3 | 5 | 7 | 3 | 5 | 7 | 3 | 5 | 7 | 3 | 5 | 7 |
| Aclust2 | 0.43 | 0.33 | 0.25 | 0.42 | 0.23 | 0.23 | 0.43 | 0.22 | 0.15 | 0.35 | 0.36 | 0.30 | 0.40 | 0.41 | 0.43 | 0.42 | 0.47 | 0.49 |
| SACOMA | 0.83 | 0.54 | 0.47 | 0.81 | 0.40 | 0.36 | 0.80 | 0.35 | 0.24 | 0.65 | 0.53 | 0.56 | 0.64 | 0.52 | 0.52 | 0.65 | 0.51 | 0.48 |
| coMethDMR | 0.99 | 0.98 | 1.00 | 0.99 | 0.98 | 1.00 | 0.99 | 0.99 | 0.99 | 0.96 | 0.98 | 1.00 | 0.96 | 0.98 | 1.00 | 0.94 | 0.98 | 1.00 |

**STable 6:** Comparison of false discovery rate (FDR) across models (Aclust2, SACOMA and coMethDMR) under varying minimum CpG thresholds (3, 5, 7) and maximum gap distances (200 bp, 500 bp, 1000 bp) for both Illumina 450K and EPICv1 arrays. These results are based on the median of 100 simulations.

| | **450K** | | | | | | | | | **EPICv1** | | | | | | | | |
|---|---|---|---|---|---|---|---|---|---|---|---|---|---|---|---|---|---|---|
| | maxGap = 200 | | | maxGap = 500 | | | maxGap = 1000 | | | maxGap = 200 | | | maxGap = 500 | | | maxGap = 1000 | | |
| | 3 | 5 | 7 | 3 | 5 | 7 | 3 | 5 | 7 | 3 | 5 | 7 | 3 | 5 | 7 | 3 | 5 | 7 |
| Aclust2 | 0 | 0 | 0 | 0 | 0 | 0 | 0 | 0 | 0 | 0.04 | 0.035 | 0.015 | 0.08 | 0.05 | 0.03 | 0.12 | 0.085 | 0.07 |
| SACOMA | 0 | 0 | 0 | 0 | 0 | 0 | 0 | 0 | 0 | 0.04 | 0.03 | 0 | 0.08 | 0.05 | 0.03 | 0.12 | 0.07 | 0.01 |
| coMethDMR | 0.03 | 0.09 | 0.09 | 0.03 | 0.11 | 0.14 | 0.03 | 0.09 | 0.13 | 0.11 | 0.14 | 0.13 | 0.15 | 0.17 | 0.17 | 0.18 | 0.18 | 0.17 |

**STable 7:** The table presents a comparison of performance metrics, genomic region characteristics, and computational resource usage across three models (SACOMA, Aclust2 and coMethDMR) applied to the EPICv1 dataset. The table summarizes the total number of detected regions, CpG density per region, unique and common region counts, peak memory consumption and execution time. The lower section presents the genomic context distribution of identified regions (Island, Shore, Shelf, and OpenSea) expressed as both counts and percentages. Counts represent the number of regions in each CpG neighborhood and the percentages (in parentheses) are their proportions out of the total number of regions identified by each model. Peak memory usage is reported in gigabytes (GB), and runtime is measured in minutes.

| **Metric** | **SACOMA** | **Aclust2** | **coMethDMR** |
|---|---|---|---|
| No. of Regions | 2385 | 1527 | 4987 |
| No. of CpGs/Region | [3, 59] | [3, 23] | [3, 62] |
| No. of Unique Regions | 1394 | 1088 | 4088 |
| Common Regions | 289 | 289 | 289 |
| Peak Memory Usage (GB) | 0.66 | 0.65 | 0.89 |
| Runtime (min) | 93.788 | 5.62 | 257.503 |
| *CpG Neighborhood* | | | |
| Island | 1126 (47.2%) | 887 (58.1%) | 1881 (37.7%) |
| N_Shelf | 34 (1.4%) | 15 (1%) | 139 (2.8%) |
| N_Shore | 324 (13.6%) | 172 (11.2%) | 674 (13.5%) |
| OpenSea | 576 (24.2%) | 286 (18.7%) | 1606 (32.2%) |
| S_Shelf | 36 (1.5%) | 13 (0.9%) | 108 (2.2%) |
| S_Shore | 289 (12.1%) | 154 (10.1%) | 579 (11.6%) |
